# Machine learning of three-dimensional protein structures to predict the functional impacts of genome variation

**DOI:** 10.1101/2023.10.27.564415

**Authors:** Kriti Shukla, Kelvin Idanwekhai, Martin Naradikian, Stephanie Ting, Stephen P. Schoenberger, Elizabeth Brunk

## Abstract

Research in the human genome sciences generates a substantial amount of genetic data for hundreds of thousands of individuals, which concomitantly increases the number of variants with unknown significance (VUS). Bioinformatic analyses can successfully reveal rare variants and variants with clear associations to disease-related phenotypes. These studies have made a significant impact on how clinical genetic screens are interpreted and how patients are stratified for treatment. There are few, if any, comparable computational methods for variants to biological activity predictions. To address this gap, we developed a machine learning method that uses protein three-dimensional structures from AlphaFold to predict how a variant will influence changes to a gene’s downstream biological pathways. We trained state-of-the-art machine learning classifiers to predict which protein regions will most likely impact transcriptional activities of two proto-oncogenes, nuclear factor erythroid 2 (NFE2)-related factor 2 (Nrf2) and c-MYC. We have identified classifiers that attain accuracies higher than 80%, which have allowed us to identify a set of key protein regions that lead to significant perturbations in c-MYC or Nrf2 transcriptional pathway activities.

**Significance:** The vast majority of mutations are either unspecified and/or their downstream biological implications are poorly understood. We have created a method that utilizes protein structure to cluster mutations from population-scale repositories to predict downstream functional impacts. The broader impacts of this approach include advanced filtering of mutations that are likely to impact genome function.

## Introduction

Massive efforts following the initial sequencing of the human genome have generated whole genome and whole exome sequencing on hundreds of thousands of individuals^1–4^. Bioinformatic analyses have greatly contributed to understanding which genomic variants are associated with clinical and disease phenotypes^5–9^. In cancer research, genome-wide association studies have been performed on the majority of common malignancies and have revealed over 450 genetic variants that are associated with increased disease risk^10–12^. These genotype-phenotype studies have been critical to understanding which genes are involved in carcinogenesis and which mutations contribute to heritable disease. However, it remains a challenge to decipher how genomic variants alter functioning on a cellular or biochemical level to influence phenotype. Understanding functional changes in cells that can explain genotype-phenotype relationships is an urgent unmet need that could change the way researchers and clinicians interpret mutation data.

With the substantial increase in genetic data, we have also seen a concomitant increase in the number of variants of unknown significance (VUS), which are genetic mutations that are not specified in their relationships to disease. Surprisingly, VUS account for 40% of identified variants in genetic screens^13,14^, which cause significant clinical burdens related to interpreting mutation profiles. For certain cancers, the likelihood that a patient will have a VUS is 91%^15^. One reason for the high percentage of VUS is not having enough phenotypic data and/or statistical power to test associations. This drastically limits the amount of clinically useful information from sequencing data. Therefore, novel methods are needed to improve our understanding of VUS to maximize the full potential of sequencing data. A first step in this direction would involve predicting which variants are most likely to induce functional changes in their respective proteins and biological pathways within cells and tissues.

The multi-layered challenge of deciphering the biological impacts of genetic variants requires integrating other functional omics datasets, together with genetic sequencing and phenotypic data, to provide clarity on how a variant influences biochemical changes within cells and tissues. Functional omics data include transcriptomics, epigenomics, proteomics, metabolomics, and molecular phenotype data, such as gene essentiality screens and drug sensitivity profiles. Large population-scale data repositories, such as Dependency Map (DepMap)^16^, Cancer Cell Line Encyclopedia (CCLE)^17–22^, and The Cancer Genome Atlas (TCGA)^23^ make these multi-omics data available for thousands of cancer cell lines as well as for patient tumors. However, understanding how to integrate and interpret the analyses of multiple disparate omics data in the context of variant effects remains unclear. Each sample may have hundreds to thousands of mutations, which makes it challenging to pinpoint which mutations are associated with which functional effects. Further, the majority of mutations across samples occur infrequently, which makes statistical analyses non-trivial.

One possible solution to these dilemmas is to cluster mutations based on how likely they are to induce protein-level changes, such as perturbations in structural conformation, binding to small molecules or ligands, binding to protein partners, and chemical reactivity. Our overarching hypothesis is that protein-level changes are more likely to occur when specific protein regions or domains, which are linked to a specific function, are mutated. To test this hypothesis, we decided to first characterize mutations based on where they occur in three-dimensional (3D) protein space. For hot-spot mutations, or regions of proteins with a higher mutational burden, we assessed the likelihood of these mutations co-occurring with significantly altered pathway activities. As a proof of principle, we predict which variants would have the highest likelihood of impacting the activities of two proto-oncogenes, nuclear factor erythroid 2 (NFE2)-related factor 2 (Nrf2) and c-MYC.

To carry out these analyses, we have developed an open-source, machine learning-based computational workflow that we refer to as <u>V</u>ariome <u>A</u>nnotations using <u>M</u>ulti-<u>O</u>mics and <u>S</u>tructural Biology (VAMOS). Using machine learning, which is a branch of artificial intelligence (AI), we uncover complex patterns and relationships in population-scale mutation datasets. We identify classifiers that attain accuracies higher than 80%, which allows us to identify key protein regions that lead to significant perturbations in c-MYC or Nrf2 transcriptional pathway activities.

## Results and Discussion

### Systematic mapping of variant effects in protein structures and downstream networks

In genetic and genomic analyses, somatic mutations or single nucleotide variants (SNV) are commonly grouped based on nucleotide positioning and frequency of occurrence within a population. While this approach has been successful in pinpointing the most common variants, it cannot determine the functional importance of the vast majority of protein-encoding variants. Furthermore, statistical associations between variants and clinical phenotypes cannot explain biological impacts on genome functioning. In recent years, structural biology approaches have been used to place protein variants in the context of their locations within three-dimensional crystallographic protein structures^24–30^. These structural genomics studies discovered that mutation clusters are enriched in specific protein domains and protein families^31–33^. However, it remains a grand challenge to systematically determine how variation at these sites in proteins leads to changes in downstream biological networks and cellular functioning.

Linking variants to their impacts on proteins as well as their impacts on downstream biological networks and cellular phenotypes is a multi-layered challenge. The different biological “layers” of determining variant effects are presented in Schema 1, which summarizes an example of each layer as well as a few common approaches that characterize changes across these layers. Layers 1 and 2 describe changes in primary amino acid sequence and alterations to protein structure and function. For example, variants may impact structural regions near active sites or allosteric binding sites, altering a protein’s catalytic function. Layer 3 translates Layer 2 to a systems-level functioning by describing changes in downstream biological networks. For example, an impaired catalytic function of one protein may induce changes in other proteins in downstream networks that are dysregulated by the altered reaction. Layer 4 translates Layer 3 to whole-cell functioning by characterizing how changes in networks lead to molecular phenotypes, such as cellular growth or sensitivity to perturbation. Finally, Layer 5 considers how all of these layers contribute to disease phenotypes present at the organism level.

**Schema 1:**
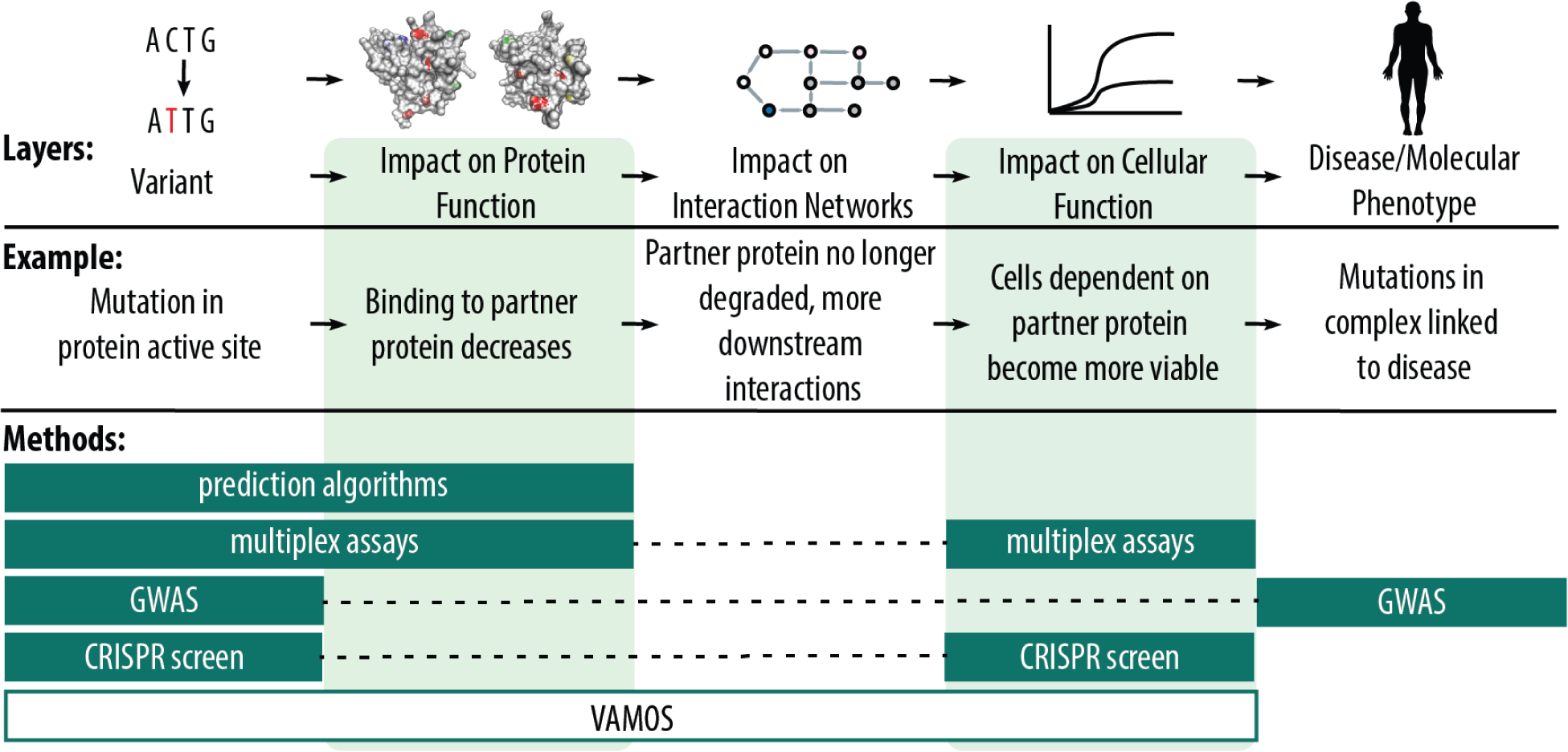
Variants have a ripple effect on function and phenotype. In recent years, genome wide association studies (GWAS) have been used to correlate primary amino acid sequence to molecular phenotypes^5,10–12,39,42^. Existing computational algorithms predict the impact of variants on protein function^103,104^, while multiplex assays of variant effects (MAVE) incorporate cellular level functional data with variant and protein functional data^31,44–46^. CRISPR screens can also provide links between primary sequence changes and cellular functioning^47–49^. However, existing methods do not address the impact on interaction networks in this ripple effect. Our proposed method, VAMOS, bridges together cellular and subcellular effects of variants.

The majority of computational chemistry methods focus on characterizing variant effects on primary amino acid sequence and protein structure and function (levels 1 and 2) using molecular dynamics, ensemble network modeling, mixed quantum mechanics methods, and free energy perturbation methods^34–38^. For computational genetics methods, changes in the primary amino acid sequences are correlated to disease phenotypes (levels 1 and 5) using methods such as genome-wide association studies (GWAS)^5,8,10–12^. In recent years, GWAS studies have also correlated primary amino acid sequences to molecular phenotypes, such as transcriptional programs^39–43^ (levels 1 and 4). Finally, experimental methods, such as multiplex assay for variant effects (MAVE)^31,44–46^, couple mutagenesis with molecular assays provide direct associations between primary amino acid sequence and impacts on protein structure/function and cellular functioning (levels 1, 2, 4). Other experimental methods, such as CRISPR screens^47–49^, can also provide links between primary sequence changes and impacts on cellular functioning (levels 1, 4) when coupled with biochemical assays, growth, transcriptomics, or other functional genomics data.

Taken together, these methods provide insights into variant effects at different levels of biological resolution. However, what is missing is a method capable of tying these different layers together to provide a more comprehensive view of variant effects. Our approach characterizes layers 1 through 4 by integrating protein structural data, functional genomics data, and molecular phenotypic data. With the help of protein structural data, we can characterize protein-coding variants, regardless of whether they occur frequently or rarely in the population. By mapping variants to their three-dimensional locations within proteins, we group them based on their proximity to other variants found in the same protein region (Fig. 1(a)). This means that variants that are far away from one another in nucleotide space can be clustered in the same groups due to the complex shapes that proteins adopt during folding. Our variant 3D clusters provide a coarse-grained, functional view of mutations that can be used to explain patterns in biological network activity, molecular phenotype data, and disease phenotype data (Fig. 1(b)).

**Figure 1.**
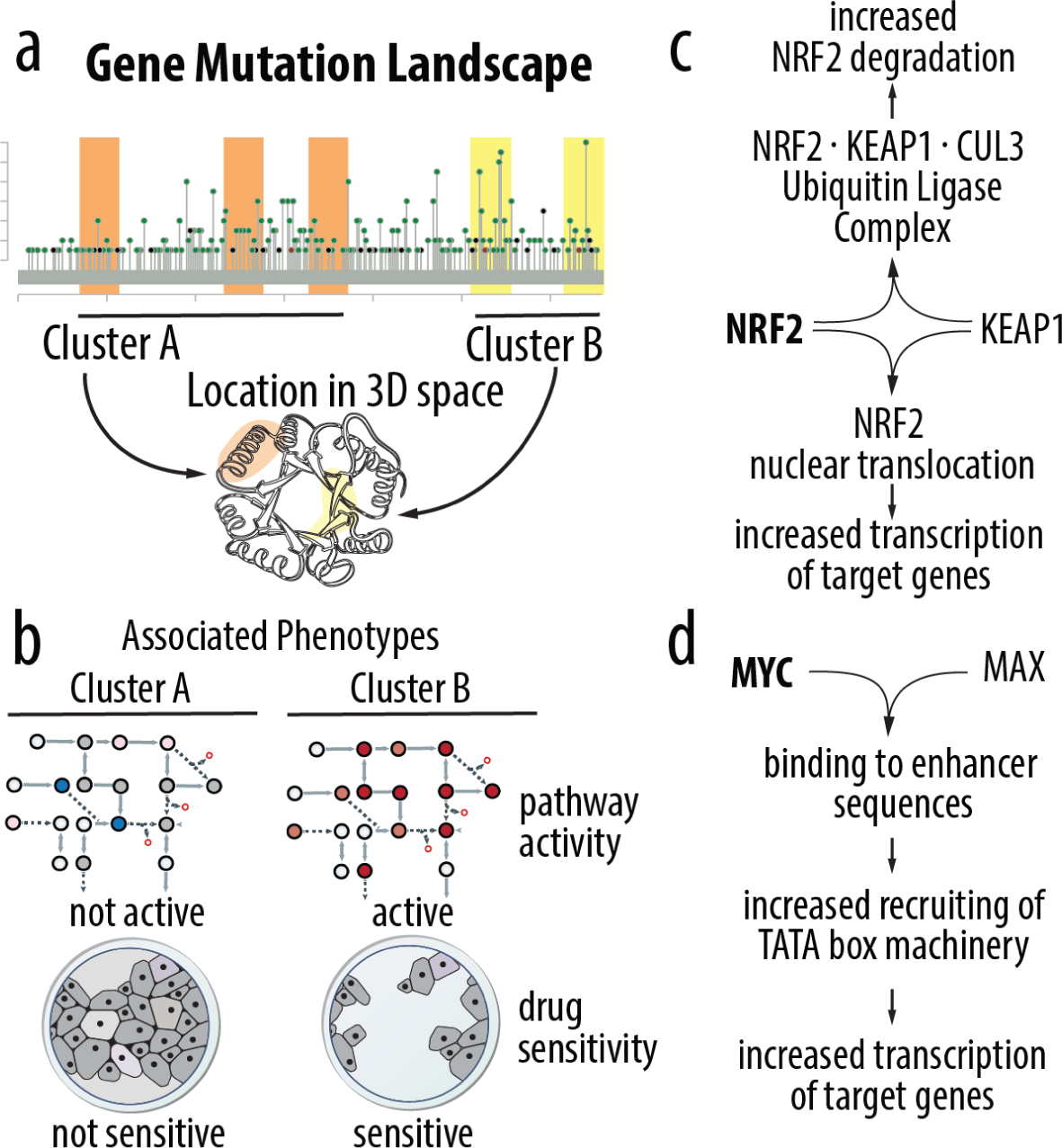
**(a)** Mutations located far from each other on the chromosome may be located close together in 3D space, forming clusters. **(b)** Different clusters may be correlated with different pathway activities, leading to different phenotypes such as drug resistance. **(c)** The NRF2 and KEAP1 interaction network demonstrates how interaction with KEAP1 can lead to increased nuclear NRF2 degradation and translocation^52,77–79^. **(d)** The c-Myc MAX interaction pathway demonstrating how c-Myc can upregulate transcription of its target genes^85,105,106^.

### Two Case Studies: NRF2 and c-MYC Transcriptional Pathway Activities

As a proof of principle, we explored two different case studies to determine whether our method could accurately predict which variants are associated with changes in biological networks and molecular phenotypes.

Our first case study characterizes the transcriptional activity of nuclear factor erythroid 2 (NFE2)-related factor 2 (Nrf2) and a subset of more than 500 genes whose transcription is regulated by Nrf2^50^. Nrf2 is responsible for regulating the cellular oxidative stress response and controls the basal and induced expression of many antioxidant response-dependent genes. Nrf2 belongs to the cap and collar (CNC) subfamily of basic region leucine zipper (bZip) transcription factors^51,52^. Nrf2 activation likely takes place via post-transcriptional mechanisms, as Nrf2 mRNA is expressed independently of inducers^53^. Under basal conditions, Nrf2 is suppressed via ubitiquination and proteasome-dependent degradation by an E3 ubiquitin ligase formed by Kelch-like erythroid cell-derived protein with CNC homology-associated protein 1 (Keap1) and other enzymes (Fig. 1(c)). In response to oxidative stress, key sensor cysteines within Keap1 are oxidized, allowing Nrf2 to escape ubitiquination, accumulate within the cell, and translocate to the nucleus^51,52^. In the nucleus, Nrf2 regulates the transcription of several genes involved in oxidative stress response. Nrf2 downstream targets are classified into three major groups: phase I and phase II drug metabolizing enzymes as well as phase III drug transporters^54^.

Our second case study characterizes the transcriptional activity of c-MYC and a subset of thousands of genes whose transcription is regulated by c-MYC^55^. c-MYC is a known oncogene that regulates many aspects of cellular growth and metabolism. The c-MYC gene encodes a helix-loop-helix leucine zipper transcription factor that can dimerize with its partner protein MAX, or form a heterodimer to transactivate gene expression^55,56^. In normal cells, c-MYC function and expression are tightly regulated and c-MYC RNA is very short-lived. In tumor cells, c-MYC is almost always overexpressed, and c-MYC is thought to contribute to the cause of at least 40% of tumors^57^. There are thousands of c-MYC target genes that span almost every important cellular function^55,57^. Notably, c-MYC is involved in the regulation of many growth-promoting signal transduction pathways^58–61^, the glucose^62–64^ and glutamine metabolic pathways^65–67^, and the regulation of genes involved in mitochondrial biogenesis^55,68–70^.

### Generating a Training Data Set To Classify Network Activities and Molecular Phenotypes

We generate training data to classify mutations based on how likely they impact network-level pathway activities and other molecular phenotypes through advanced data integration. Our integration approach consists of three main stages, as shown in Figure 2. In the first stage (Fig 2, Box 1), we use variant information from two large-scale cancer population-scale data repositories, the Cancer Genome Atlas (TCGA)^23^ and the Cancer Cell Line Encyclopedia (CCLE), which is also referred to as the Dependency Map (DepMap)^16^. TCGA provides population-scale mutation data for over 20,000 primary cancer and matched normal samples spanning 33 cancer types, taken from cancer patient tumor biopsies with over 1.6 million protein-coding variants across 22,444 genes. DepMap/CCLE provides population data for over 2,000 cancer cell line samples with over 1.2 million protein-coding variants across 19,537 genes. Second, we filter all variants based on whether experimental crystallographic information or computationally predicted structural information was available. For computationally predicted structures, we consider Alphafold^71^ structures with structural prediction confidence scores higher than 70% (Fig 2, Box 2). Our filtering finds 17,062 genes with available structural data, where 896,629 mutations have experimental coverage or highly confident structural predictions. Third, we perform density-based scanning to cluster all mutations for each gene based on their locations within three-dimensional protein structures. Variant clusters represent a conglomerate of mutations observed across samples within the DepMap/CCLE or TCGA databases (Fig 2, Box 3).

**Figure 2.**
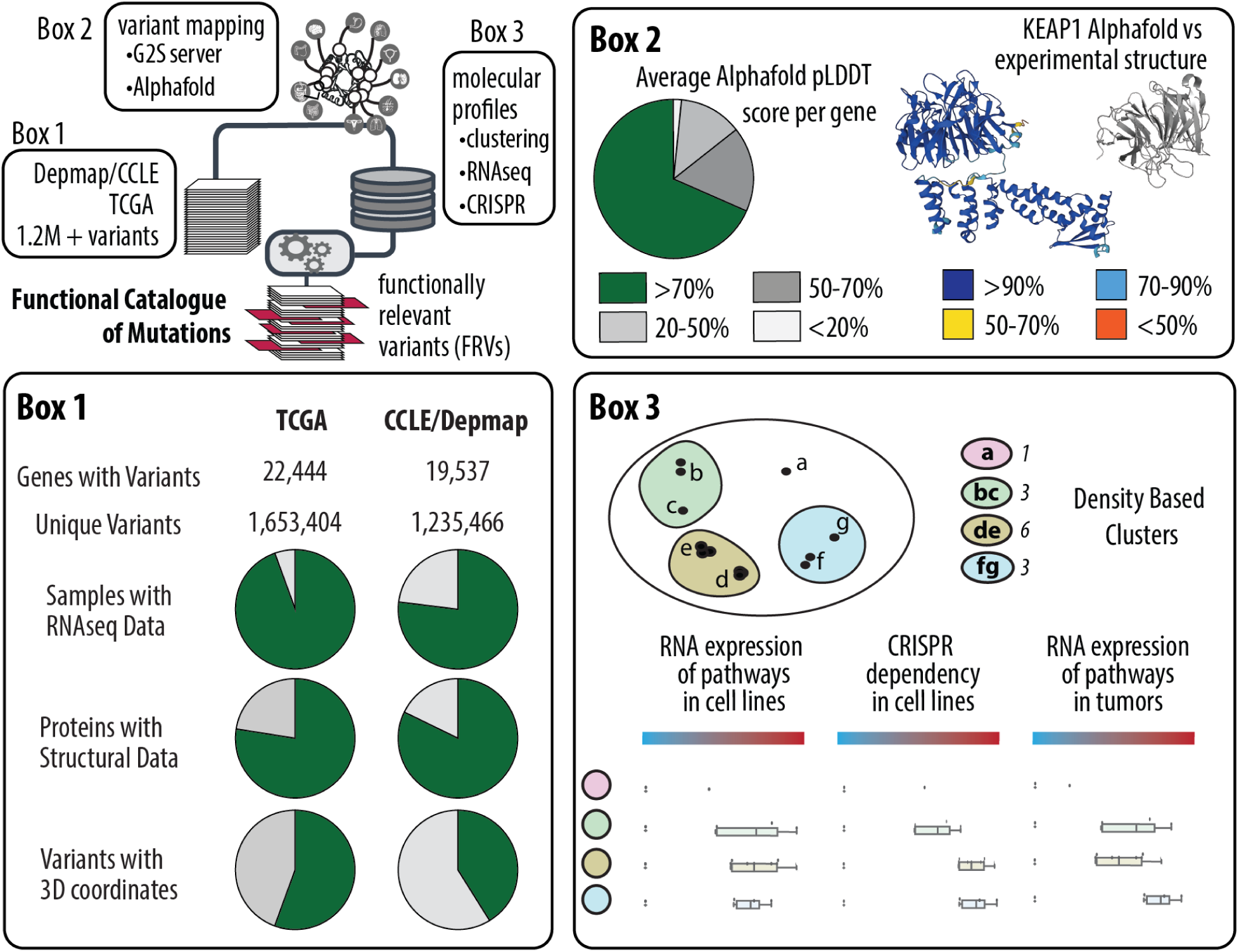
An overview of the proposed pipeline to extract functionally relevant variants from VUSs. Variants are taken from DepMap/CCLE and TCGA. The variants are mapped to 3D protein structures in the PDB using the G2S server, and to AI-predicted Alphafold structures. Clustering is performed on the structural information, and after machine learning, functionally relevant variants can be filtered out. **Box 1** shows a summary of available genomics and proteomics data for variants from DepMap/CCLE and TCGA. Structural data is from Alphafold and PDB structures. **Box 2** shows an overview of the average pLDDT (predicted local difference test) scores for each protein structure in Alphafold. pLDDT scores >70% indicate high confidence of prediction. A comparison of the KEAP1 structure as predicted by Alphafold and as experimentally determined is also shown. **Box 3** shows a comparison of density based clustering methods vs hierarchical clustering methods on the same example dataset.

To classify the functional impacts of this set of mutations, we integrate molecular phenotype data as outcome variables (or response variables). First, we consider transcriptional regulation and pathway activities of NRF2 and c-MYC. Transcriptional data, or bulk RNA sequencing data, is available for 1,210 samples in DepMap/CCLE and 8,635 samples in TCGA. Second, we consider genome-wide CRISPR-mediated genetic knockdown screens, which have been generated for 1,086 samples in DepMap/CCLE. We integrate cell line sensitivity profiles for these samples where NRF2 or c-MYC has been genetically knocked down.

Using this approach, we collect training data for four different model predictions: (i) variants that associate with increased gene expression for genes that are transcriptionally regulated by NRF2; (ii) variants that associate with increased gene expression for genes that are transcriptionally regulated by c-MYC; (iii) variants that associate with increased sensitivity to NRF2 knockdown; (iv) variants that associate with increased sensitivity to c-MYC knockdown.

### Training and Optimizing Machine Learning Models

Once the training data has been built, it is used to guide learning of which mutations are more significantly associated with the four molecular phenotypes and to pinpoint which protein regions are more likely to impact downstream biological activities. Our overall approach to identifying functionally relevant mutations is two-fold. First, we divide our dataset into training (66.6%, 411,844 mutations across 5,757 samples from DepMap/CCLE with transcriptomics data) and testing (33.3%, 205,922 mutations across 2,878 samples) subsets. Second, we apply two types of machine learning algorithms to classify mutations that are associated with discrete molecular responses. We apply a random forest classifier. One of the main benefits of this machine learning algorithm is that it is capable of higher-order interactions between features and can often achieve higher accuracies when compared to other approaches^72,73^. Here, the features that we use as input to our machine learning models are the atomic positions (cartesian coordinates) of each mutation, its respective cluster which was determined through a density-based scanning method, and information about the cluster such as the average distance to centroid and number of members. While these are a relatively small number of features, our models are capable of predicting which mutations are significantly associated with specific molecular phenotypes (AUC = ∼.84-.89, Fig. 3(b)). This means that the atomic positions of amino acids in proteins provide significant capacity to learn patterns and recognize the regions of proteins most likely to influence downstream pathway behavior and cellular phenotypes.

**Figure 3.**
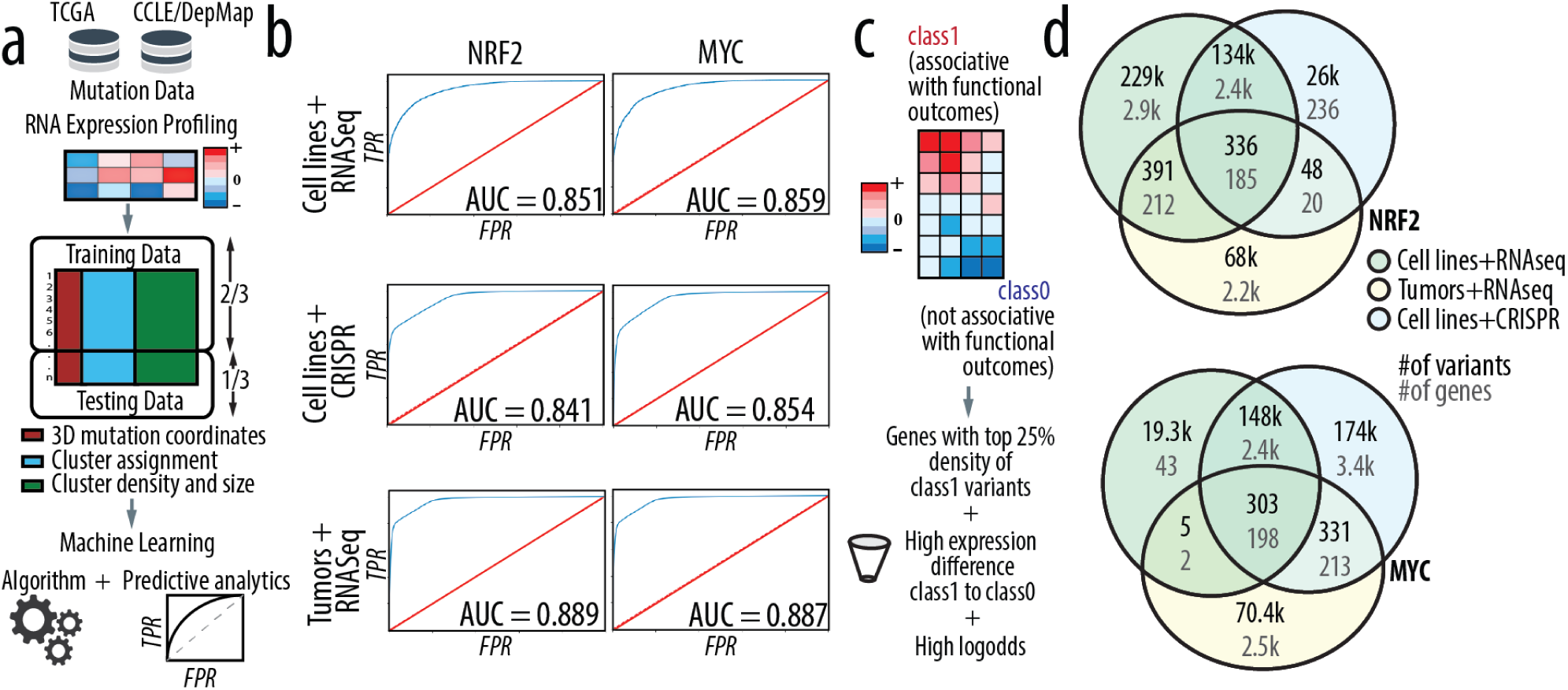
**(a)** An overview of the Machine Learning pipeline to identify functionally relevant variants. The dataset is comprised of mutation and expression data. ⅔ of the dataset is used as training data, and ⅓ as testing data, with 5-fold cross-validation. The features include structural information, clustering assignments, and cluster density and size. Predictive analytics are performed on the algorithm to determine quality. **(b)** ROC plots for datasets balanced by oversampling. Here, all combinations of DepMap/CCLE and TCGA with RNAseq expression data and CRISPR sensitivity data for NRF2 and c-MYC targets are shown. **(c)** The machine learning algorithm assigns variants as class 1 (functionally relevant) or class 0 (not functionally relevant). Variants are then additionally filtered. Only genes in the upper 25% of class 1 variant density are chosen. Genes also need to be over one standard deviation above the mean in terms of normalized expression differences between class1 and class0 variants. Finally, genes need to be one standard deviation above the mean in the log odds of a variant being assigned as class1 given that it is in a specific cluster and that it has a high expression score. **(d)** Variants with structural data present in Depmap/CCLE and TCGA using RNAseq expression data and CRISPR data after all machine learning and filtering steps have been performed.

### Data Quality, Ranking, and Prediction Consistency

Our initial predictions indicated that a large number of variants were significantly associated with at least one of the molecular response variables. We refer to associating mutations as “Class 1,” as compared to “Class 0,” which do not have a significant association with a response variable. When comparing the number of Class 1 mutations for each response variable, we find that between 100K-115K variants associate with transcription pathway activities for both Nrf2 and c-MYC molecular phenotypes. We find a large number of mutations that are consistent across cell line model data and tumor data from TCGA (4.6K and 839 variants for Nrf2 and c-MYC, respectively). We also find a large number of variants that are associated with CRISPR-mediated knockdown sensitivity of Nrf2 and c-MYC (60.5K and 424K variants, respectively). Variants with the most consistency across the three datasets for each molecular phenotype were significantly high, in the range of 673 for c-MYC and 893 for Nrf2. Variants that are consistently associated with each of the three response variables signify the highest confidence mutations that most likely influence cellular functioning with respect to Nrf2 or c-Myc transcriptional activities. Of the variants that are labeled as Class 1, we find that the top five most enriched protein domains include immunoglobulins, zinc finger, cytochrome complex, P-loop containing NTP hydrolase, phosphotransferase, and Von Willebrand factor domains (Fig. SI 2).

We also filter mutations based on other confidence scoring metrics, in the context of cluster behavior and similarities of associations within clusters (Fig. 3 (c)). First, we rank genes with the highest density mutation clusters, which signifies regions of a protein with the highest amount of mutational burden. Second, we compute the GSEA expression pathway scores between the mutations of a given cluster and mutations that are classified as class 0. Third, we compute the log odds for mutations within a cluster and rank all clusters based on log odds; the clusters with the highest log odds represent clusters with the highest association with upregulated or downregulated pathway activities or sensitivities to genetic knockdown. Taken together, these filtering steps generate a subset of genes with the most dense mutation clusters that are significantly associated with the response variable.

Post filtering, we find 727 variants for Nrf2 and 308 for c-MYC that are consistent across cell line model data and tumor data from TCGA. There are still a large number of variants that are associated with CRISPR-mediated knockdown sensitivity of Nrf2 and c-MYC (26k and 174k variants, respectively). We find 336 variants for Nrf2 and 303 variants for c-MYC that are consistently associated with all three response variables.

### Mutations Associated with Distinct Nrf2 Molecular Phenotypes

Understanding which mutations attenuate or upregulate NRF2 transcriptional activities would greatly benefit clinical practice, as it has been shown that mutations in certain genes increase Nrf2 activity and determine patient response to certain chemotherapeutics^74^. We use the Nrf2 transcriptional pathway as a proxy for Nrf2 activity and define its pathway using the hallmark pathway dataset generated using Gene Set Enrichment Analysis (GSEA)^75^, stored in MSigDB^76^. The Nrf2 pathway contains over 500 genes that were detected to have significant gene expression changes in response to Nrf2 knockdown. These genes are thought to be direct Nrf2 targets or be impacted by Nrf2 regulation. Using this set of genes, we perform GSEA to score RNA sequencing data for 1,180 cell line models from DepMap/CCLE and 8,167 tumor samples from TCGA. Using the GSEA scores for each dataset, we rank samples as “Nrf2 High” if they fall within the top 25 percentile and “Nrf2 low” otherwise. Our machine learning models use these scores as response variables to predict which mutations are consistently classified as associated (class 1) versus not-associated (class 0) with “Nrf2 High” or “Nrf2 Low” phenotypes.

Using this approach, we find several examples that are consistent with previous studies. For example, Keap1 (Kelch-like ECH-associated protein 1) is a protein that acts as an adaptor subunit of the Cullin 3-based E3 ubiquitin ligase degradation system^51,52^. Keap1 is known to regulate the transcriptional activity of Nrf2 during cellular stress by influencing its degradation^51,52^. Our findings are consistent with previous findings^77–79^ that report key mutations in Keap1 prevent binding of Nrf2, leading to constitutive Nrf2 transcriptional activity as a result of increased Nrf2 protein abundance. We find that mutations in the same region of KEAP1 are significantly associated with increased Nrf2 pathway activity in both cell line models and tumors and that some of these mutations are also significantly associated with increased sensitivity to CRISPR-mediated Nrf2 knockdown (Fig 4(a), Fig. 5(a)). When we compare Nrf2 pathway activity and sensitivity to Nrf2 knockdown in Class 1, Class 0, and mutations across other cell lines, we see significant differences (p = 2.51 x 10-14), suggesting that these mutations represent potential biomarkers for disruption of Nrf2 pathway activity as well as for cellular dependencies on Nrf2 activity.

**Figure 4.**
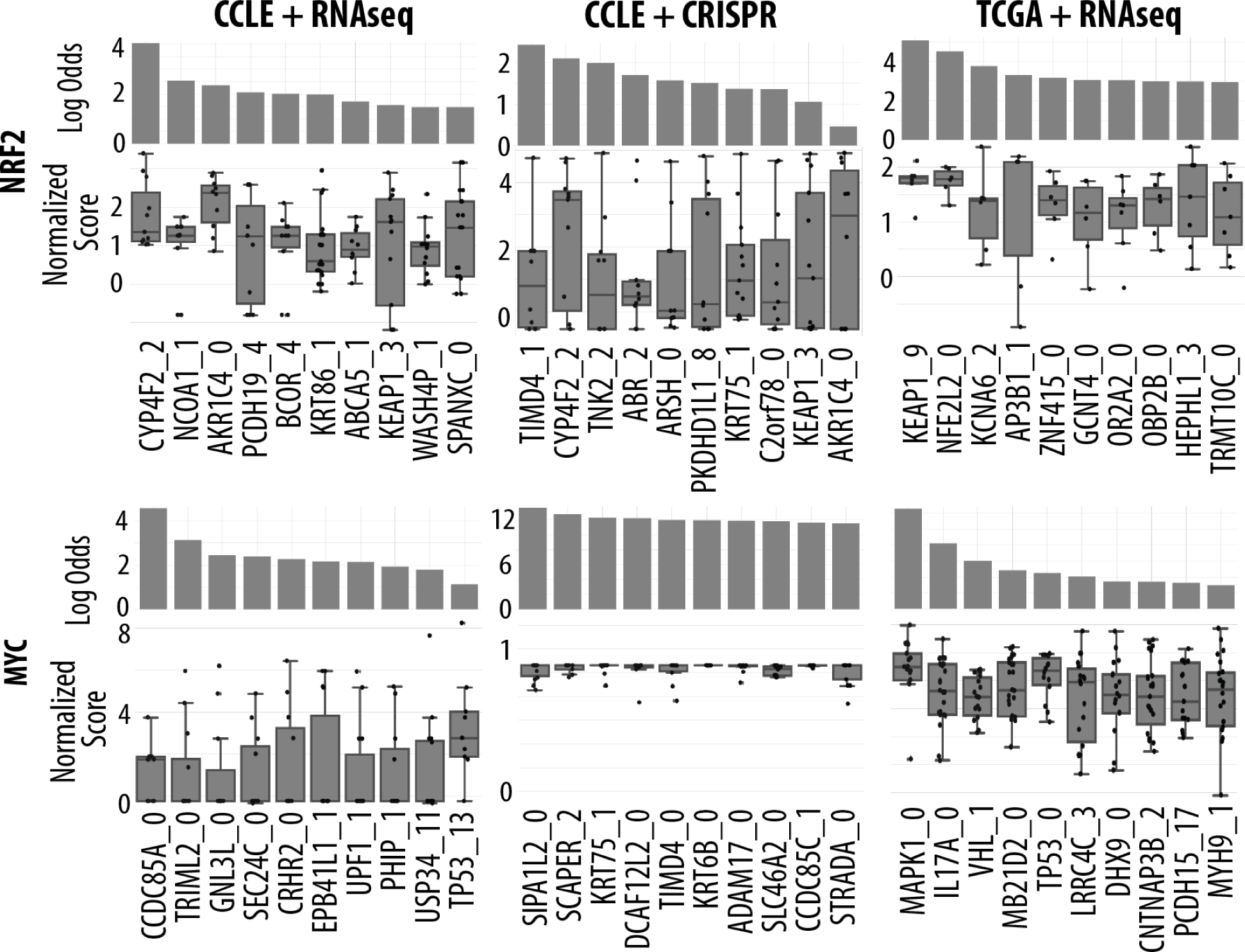
Top 10 clusters with the highest log odds scores after filtering for Nrf2 and c-MYC. These clusters are shown for all combinations of DepMap/CCLE and TCGA with GSEA pathway data and CRISPR sensitivity data for NRF2 and c-MYC targets, after filtering was performed.

**Figure 5.**
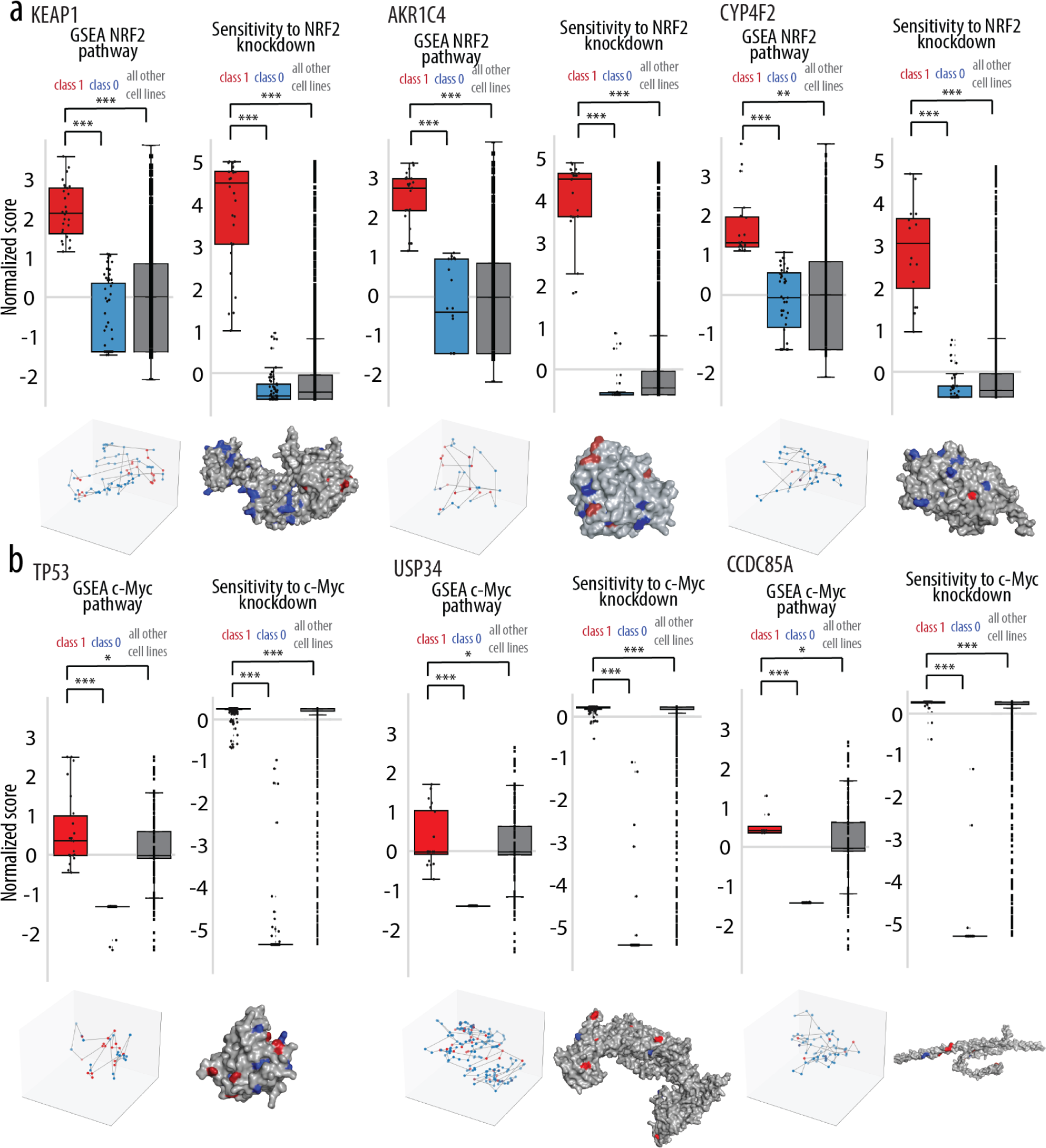
**(a)** Selected proteins that passed filtering criteria for the VAMOS workflow with NRF2 as the target. Normalized GSEA scores and Sensitivity to NRF2 knockdown is shown between class1 mutations, class0 mutations, and all cell lines present in DepMap/CCLE **(b)** Selected proteins that passed filtering criteria for the VAMOS workflow with c-MYC as the target. Normalized GSEA NRF2 pathway scores and Sensitivity to NRF2 knockdown is shown between class1 mutations, class0 mutations, and all cell lines present in DepMap/CCLE

We also observe a highly dense mutation hotspot in the Aldo-Keto Reductase 1C4 (AKR1C4) gene (5 variants in a 1.51 Angstrom sphere). AKR1C4 is part of the Aldo-Keto Reductase family of enzymes, which play a key role in the metabolism of steroid hormones by catalyzing NADPH-dependent aldehyde and ketone reduction reactions^80^. Other enzymes in the AKR family have been shown to be induced by Nrf2 mediated indirectly via an antioxidant response element (ARE)^81,82^. Based on the Nrf2 consensus sequence, AKR1C4 contains 2 AREs and is likely regulated by Nrf2^81^. Similar to Keap1, we find that Class 1 mutations in AKR1C4 are significantly higher in Nrf2 pathway activities compared to Class 0 and mutations from other cell lines. Moreover, we see a consistent pattern demonstrating that cell lines with Class 1 mutations are more likely to be sensitive to Nrf2 knockdown (Fig. 4(a), Fig. 4(b)).

Another example of a gene with mutations highly associated with Nrf2 pathway activities is CYP2C4. CYP2C4 is a member of the Cytochrome P450 (CYP) system. The CYP system plays a key role in the metabolism of drugs and is of great interest to practitioners^83^. Previously, CYP2C4 has been shown to be a biomarker for liver cancer. Overexpression of CYP2C4 has been shown to repress the expression of Nrf2 and other genes in the Nrf2 signaling pathway^84^. In our analyses, 19 Class 1 mutations in CYP2C4 are found to be significantly associated with upregulated Nrf2 pathway expression in cell line models. Cell lines with these mutations are also more likely to be sensitive to Nrf2 knockdown (Fig. 4(a), Fig. 4(b)).

### Mutations Associated with Distinct c-MYC Molecular Phenotypes

Understanding which mutations attenuate or upregulate c-MYC transcriptional activities would greatly benefit clinical practice, as c-MYC is a known oncogene. c-MYC modulates over 15% of the known human transcriptome, and the deregulation of c-MYC occurs in over 70% of human cancers^85^. Therefore, c-MYC is a promising therapeutic target molecule. Similar to our approach for Nrf2, we use the c-MYC transcriptional pathway as a proxy for c-MYC activity and define its pathway using the hallmark pathway dataset generated using Gene Set Enrichment Analysis (GSEA)^86^, stored in MSigDB^76^. c-MYC regulates the transcription of thousands of genes^55^. Using this set of genes, we perform GSEA to score RNA sequencing data for 1,100 cell line models from DepMap/CCLE and 8,167 tumor samples from TCGA. Similar to our approach for Nrf2, using the GSEA scores for each dataset, we rank samples as “c-MYC High” if they fall within the top 25 percentile and “c-MYC low” if they fall within the lowest 25 percentile of c-MYC pathway expression within the population. Our machine learning models use these scores as response variables to predict which mutations are consistently classified as associated (class 1) versus not-associated (class 0) with “c-MYC High” or “c-MYC Low” phenotypes.

Using this approach, we find several examples for c-MYC that are consistent with previous studies. For example, tumor protein 53 (TP53) acts as a tumor suppressor by keeping cells from proliferating^87^. TP53 suppresses c-MYC through a combination of transcriptional inactivation and RNA-mediated repression^88^, while c-MYC induces the expression of TP53^88^. Our findings are consistent with previous findings^89–91^ that report that c-MYC is an important target of mutant TP53. We find that mutations in the same region of TP53 are significantly associated with increased c-MYC pathway activity in both cell line models and tumors (Fig 4(b), Fig. 5(b)). When we compare c-MYC pathway activity and sensitivity to c-MYC knockdown in Class 1, Class 0, and mutations across other cell lines, we see significant differences (p = 2.39 x 10-15), suggesting that these mutations represent potential biomarkers for disruption of c-MYC pathway activity as well as for cellular dependencies on c-MYC activity.

We also see similar trends in other genes where the biological connection to c-MYC is less clear. For example, we observe that the Ubitiquin Specific Peptidase 34 (USP34) gene also has several mutations highly associated with c-MYC pathway activities. USP34 is a deubiquitylating enzyme that has been shown to localize at DNA damage sites and promote cellular responses to DNA damage^92^. Additionally, USP34 is known to regulate axin stability^93^. When localized to the nucleus, AXIN2 co-occupies β-catenin/T-cell factor complexes at the c-MYC promoter region and directly represses c-MYC gene expression^94^. Similar to TP53, we find that Class 1 mutations in USP34 are significantly higher in c-MYC pathway activities compared to Class 0 and mutations from other cell lines. We additionally see a consistent pattern demonstrating that cell lines with Class 1 mutations are more likely to be sensitive to c-MYC knockdown (Fig. 4(b), Fig. 5(b)).

Another example of a gene with mutations highly associated with c-MYC pathway activities is Coiled-Coil Domain Containing 85A (CCDC85A). CCDC85A is a member of the delta-interacting protein A (DIPA) family. Other members in this family, such as CCDC85B and CCDC85C, are regulated by TP53 and have been shown to regulate β-catenin which regulates c-MYC gene expression^95^. In our analyses, 8 Class 1 mutations in CCDC85A are found to be significantly associated with upregulated c-MYC pathway expression in cell line models. Cell lines with these mutations are also more likely to be sensitive to c-MYC knockdown (Fig. 4(b), Fig. 5(b)).

### Precision Medicine Applications: NRF2 Adjuvant Therapies

Cellular dependencies on oncogenes like Nrf2 represent opportunities for targeting vulnerable cancer cells that require Nrf2 to proliferate. One approach is to develop combination therapies that involve standard chemotherapies as well as adjuvant therapies that target a more specific cellular vulnerability. However, determining which patients should be given adjuvant treatment requires representative biomarkers. In this study, we assess large population-scale mutation landscapes to determine which mutations were most likely associated with specific molecular pathway activities and cell dependencies related to Nrf2. We consider our predicted set of variants to be potential biomarkers that signify higher Nrf2 pathway activities and tumor cells with higher dependencies on Nrf2. Our positive control, A549 lung cancer cells, have a mutation in the binding site of Keap1 that is known to prevent degradation of Nrf2, thereby leading to Nrf2 being constitutively active. In contrast, our negative control, NCIH2444 lung cancer cells, did not have mutations in any of the protein regions predictive for increased Nrf2 activity. As shown in Fig 6(d), Nrf2 pathway activities and Nrf2 dependency scores are significantly different between the positive and negative controls.

**Figure 6.**
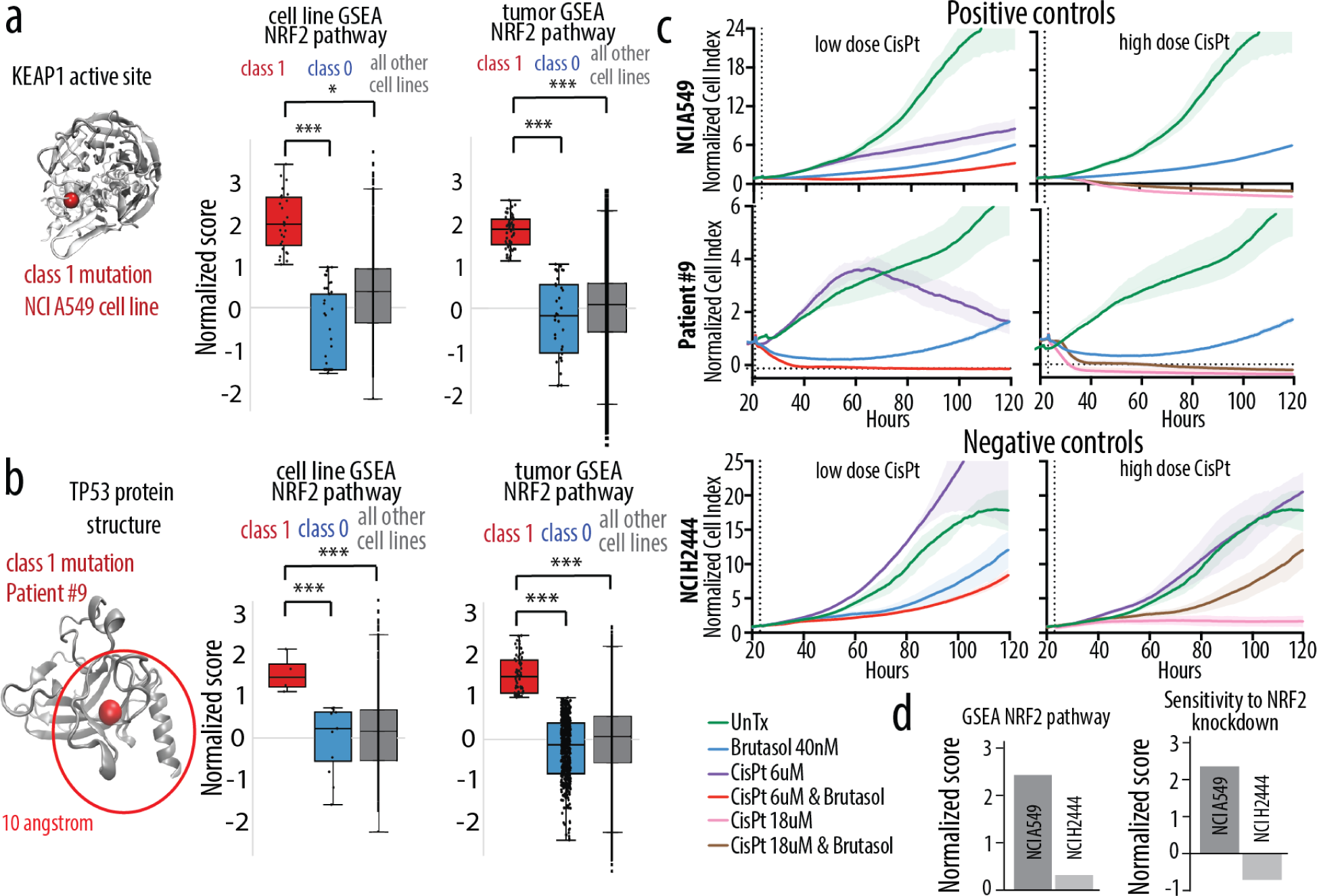
**(a)** The class1 mutation identified by VAMOS in KEAP1 in the NCIA549 cell line. Normalized scores for the GSEA NRF2 pathway are shown between class1 mutations, class0 mutations, and all cell lines present in DepMap/CCLE for the cluster this mutation is in, using TCGA and Depmap/CCLE data. **(b)** The class1 mutation identified by VAMOS in TP53 from Patient #9’s mutational landscape. Normalized scores for the GSEA NRF2 pathway are shown between class1 mutations, class0 mutations, and all cell lines present in DepMap/CCLE for the cluster this mutation is in, using TCGA and Depmap/CCLE data. **(c)** RTCA growth curves demonstrating the effect of different chemotherapies on the positive controls: NCIA549 cell line and Patient #9, and the negative control: the NCIH2444 cell line. Here the effects of UnTx, Brustasol, Cisplatin (6 and 18 μM), and Brustasol+Cisplatin(6 and 18 μM) on cell growth was monitored for 120 hours. **(d)** Bar graphs showing the normalized scores for GSEA NRF2 pathways and the sensitivity to NRF2 knockdown for the NCIA549 cell line and the NCIH2444 cell line

Our machine learning model predicted that a patient-derived cell line model (referred to as patient #9) was likely to have activated Nrf2 programming and potentially be sensitized to Nrf2 inhibition. One particular mutation in TP53 stood out as particularly predictive for increased Nrf2 pathway activities and increased sensitivities to Nrf2 knockdown (Fig 6(b)). To apply this knowledge, we perform real-time continuous growth monitoring analysis (RTCA) in the presence of a standard chemotherapy (Cisplatinum) as well as with a Nrf2 adjuvant (Brutasol). We evaluated the impact of each single drug as well as drug combinations on our positive control, A549 cells, patient #9 and our negative control, NCIH2444. Both samples our positive control and patient #9 cells have a subset of Class 1 mutations, indicating higher Nrf2 pathway levels and higher dependencies on Nrf2 (Fig. 6(a), Fig. 6(b)). Our experiments consisted of monitoring the continuous growth (a measurement every 15 minutes) over the course of 120 hours for 8 different drug treatment conditions. We compared untreated cells to those treated with: (i) low dose of cisplatin; (ii) high dose of cisplatin; (iii) Nrf2 inhibitor; (vi) low dose of cisplatin + Nrf2 inhibitor; (v) high dose of cisplatin + Nrf2 inhibitor. We found that for both samples, the combination of cisplatin and Nrf2 inhibitor significantly decreased the dose of cisplatin while maintaining effective cell killing (Fig. 6(c)). For our negative control, we find that the combination of cisplatin and Nrf2 inhibitor does not have as significant of an effect on cell killing as in the A549 cells or patient cells (Fig. 6(c)).

### Conclusion

We developed a bioinformatics approach for predicting variants that have functional impacts on downstream transcriptional networks and cellular dependencies/sensitivities from protein structural data. The ability to rank variants based on the likelihood that they are involved or associated with biological functions and cellular phenotypes will make it possible to prioritize mutations for further studies on clinical impact as well as to explain the biological impacts of well-known clinical variants. Doing so will decrease the number of variants of unknown significance and provide detailed information on not only variant clinical effects but also on variant biological effects. The novelty of this approach is that we use protein structural information to engineer explainable, biologically informed features that guide the learning of impactful variants. From our study, we show that three-dimensional locations within proteins are sufficient in providing a highly predictive algorithm, with AUC between 0.84 and 0.89. We applied this method on two case studies to predict variants that are significantly associated with transcriptional activities of Nrf2 and c-MYC. For each gene, we used three different response variables: (i) GSEA-based transcriptional pathway scores using expression data from 1,840 cell lines in DepMap/CCLE; (ii) GSEA-based transcriptional pathway scores using data from 8,635 tumors in TCGA; (iii) CRISPR loss-of-function screens for 1,086 cell lines in DepMap/CCLE. Classifiers’ performance gave a range of variants that were significantly associated with each of these response variables. Filtering and ranking of variants led to a candidate subset that is consistently significant in their association to all three response variables for each gene. This method requires population-scale data in order to have a large enough sampling size of variants that form three-dimensional clusters in proteins. This method performs optimally when response variables clearly define cellular phenotypes. For example, for transcription factors, like Nrf2, variants in proteins that directly interact with Nrf2 are likely to impact Nrf2 activity, and this impact is demonstrated in the RNA abundance levels of Nrf2 target genes. We applied this analysis to test its translation to precision medicine by selecting samples that would be more or less vulnerable to Nrf2 adjuvant therapies. This analysis demonstrated the power of machine learning to predict which patients will respond more effectively to Nrf2 adjuvant therapies. The concordance of predictions using multiple datasets demonstrates that trained classifiers can be used to identify protein regions and the molecular activities that are directly impacted if they are mutated. Future work will be focused on validating our machine-learning approach as a tool to accelerate the functional annotation of variants and on investigating the biological connections between disruption of key protein regions and dysregulated networks and cellular functioning.

## Methods

### Data Retrieval

We obtained mutation and CRISPR dependency data from Depmap/CCLE Version 22Q2 and TCGA (version 2). Depmap/CCLE contains information for 1,653,404 variants across 22,444 genes, and TCGA contains information for 1,235,466 variants across 19,537 genes. The CRISPR Dependency data is in the form of probability scores representing the likelihood of cell death when the gene is knocked out. Mutation and transcriptomics data for patient-derived cell line, patient #9, was generated prior to this study

### Gene Set Enrichment Analysis

We ran GSEA using RNA expression data through Gene Pattern Notebook^75^ using hallmark gene sets taken from MSigDB^76^. The two gene sets that were used in this study were Nrf2^96^ and Myc^97^. Default parameters were used to run single sample GSEA^86,98^.

### Cell Culture and Real-time Continuous Analysis / Growth Monitoring Assay

Patient #9 (La Jolla Institute for Immunology, LJI) NCIH244 (ATCC) and A549 cells (ATCC) were grown in RPMI media (Gibco) supplemented with 10% heat-inactivated fetal bovine serum (Gibco) in a humidified incubator with 5% CO2. Cells were harvested with 0.25% Trypsin in DPBS (Gibco). Viable cells were counted using a Countess 3 (Invitrogen) counter and trypan blue (Invitrogen, T10282). An xCELLigence real-time cell analyser (Agilent) was used to monitor cell activity over time, recording sweeps every 15 minutes using RTCA Software Pro 2.6.0. The Agilent SP station was placed in a humidified CO2 incubator. Background readings were recorded using 80μL media alone in a 96 Well PET e-plate. Cells were seeded at a concentration of 20,000 cells per well in 100μL of media. After 48 hours of growth, appropriate concentrations of drugs were added in a total volume of 20μL (or dimethysulfoxide as the control). The following drugs were assayed: Cisplatinum (XXX), Brutasol (XXX), which were dissolved in a stock solution of 10mM in DMSO. Cell index was assayed over 96 hours after addition of drug. To avoid evaporation induced edge-effects, the outer wells of the 96-well plate were filled with DPBS. Cell index was normalized.

### Mapping to the Protein Data Bank and Alphafold

We mapped the variants to the Protein Data Bank on a per residue level using the mmtf-Python workflow developed by Peter Rose^99^ and the G2S server^100^, and to Alphafold v3 using the gemmi Python package^101^. We obtained the xyz coordinates for each mutation. For mutations mapped to Alphafold, mutations in disordered regions or with a plDDT confidence score lower than 70% were removed. pLDDT scores were based on the lDDT-C alpha matrix and obtained directly from Alphafold. From Depmap/CCLE, 511,076 variants across 16,092 genes had structural data that met the above criteria. From TCGA, 918,614 variants across 16,074 genes had structural data that met the above criteria.

### Density-Based Clustering of Mutations

We performed density-based clustering on the xyz mutation coordinates using the scikit-dbscan^102^ package on a per-gene basis. The selection of the epsilon parameter was automated using the kneed package, allowing for rapid clustering across all genes without human interference.

### Machine Learning

We used the scikit-learn^102^ package to perform all machine learning. We created the training dataset by assigning mutations a preliminary classification of 0 (not of interest with respect to target activity) or 1 (of interest with respect to target activity) based on their expression and sensitivity scores. Any cell line with a score in the top 25% was assigned as class 1, and any cell line with a score in the bottom 25% was assigned as class 0.

We selected the xyz coordinate information, density-based clusters, distance of each mutation to the centroid of the cluster, and the number of members in each cluster as our features for machine learning. The data was balanced between classes via oversampling. One-third of the data at random was used as the test set and two-thirds as the training set, with 5-fold cross-validation. Accuracy, Recall, Precision, F1 scores, AUC, and MCC scores were calculated for all datasets.

### Statistical Analysis and Filtering

We calculated the log odds that a class 1 mutation would be in a cluster using the formula:

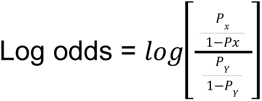

where Px is the probability of being a class1 mutation and Py is the probability of being a class 0 mutation.

We additionally filtered for clusters that contained the top 25% of density of class 1 mutations, and that were one standard deviation above the mean in terms of the difference in normalized expression scores between class 1 and class 0 mutations.

We used the scipy-stats package to perform Wilcoxon rank-sum tests to determine significance of all our analyses.

### Data availability

The workflow for this project has been compiled into a series of user-friendly Jupyter notebooks available at:

**Table 1:** Availability of gene and structural data across Uniprot, Depmap/CCLE, Alphafold and the PDB.

|  | Total Genes in Human Genome (in Uniprot) | Genes with mutations in DepMap/CCLE | Genes with Alphafold predicted structures | Genes with crystallographic /NMR structures |
| --- | --- | --- | --- | --- |
| Total genes | 20,285 | 19,537 | 23,391 | 7,788 |
| Genes with active site data | 3,534 | 3,299 | 2,017 | 1,001 |
| Genes with binding site data | 26,028 | 19,608 | 4,022 | 1,992 |
| Total number of amino acid DepMap mutations | 1,235,466 | 1,235,466 | 548,075 | 60,956 |
| Total number of amino acid DepMap mutations that are active site residues | 3,299 | 1,086 | 113 | 69 |
| Total number of amino acid DepMap mutations that are binding site residues | 19,608 | 3,254 | 737 | 100 |

**Table 2:** Scoring metrics for the machine learning algorithm applied to the training data set.

|  | Nrf2 |  |  | c-MYC |  |  |
| --- | --- | --- | --- | --- | --- | --- |
|  | Cell lines + RNAseq | Cell lines + CRISPR | Tumors + RNAseq | Cell lines + RNAseq | Cell lines + CRISPR | Tumors + RNAseq |
| Accuracy | 0.972<br>$\pm 4.27 \times 10^{-5}$ | 0.987<br>$\pm 1.58 \times 10^{-4}$ | 0.982<br>$\pm 5.04 \times 10^{-4}$ | 0.990<br>$\pm 1.34 \times 10^{-4}$ | 0.954<br>$\pm 0.001$ | 0.974<br>$\pm 0.001$ |
| Precision | 0.952<br>$\pm 0.001$ | 0.976<br>$\pm 1.82 \times 10^{-4}$ | 0.970<br>$\pm 5.05 \times 10^{-4}$ | 0.981<br>$\pm 2.23 \times 10^{-4}$ | 0.977<br>$\pm 0.002$ | 0.955<br>$\pm 0.001$ |
| Recall | 0.995<br>$\pm 0.001$ | 0.998<br>$\pm 1.93 \times 10^{-4}$ | 0.997<br>$\pm 5.03 \times 10^{-4}$ | 0.999<br>$\pm 6.13 \times 10^{-5}$ | 0.932<br>$\pm 0.002$ | 0.995<br>$\pm 0.001$ |
| F1 score | 0.973<br>$\pm 3.91 \times 10^{-5}$ | 0.987<br>$\pm 1.53 \times 10^{-4}$ | 0.983<br>$\pm 4.85 \times 10^{-4}$ | 0.990<br>$\pm 1.36 \times 10^{-4}$ | 0.954<br>$\pm 0.001$ | 0.975<br>$\pm 0.001$ |
| MCC score | 0.945<br>$\pm 8.02 \times 10^{-5}$ | 0.973<br>$\pm 3.17 \times 10^{-4}$ | 0.865<br>$\pm 0.001$ | 0.981<br>$\pm 2.64 \times 10^{-4}$ | 0.913<br>$\pm 0.002$ | 0.999<br>$\pm 1.96 \times 10^{-6}$ |

\*standard deviation values are from 5-fold cross validation
\*balanced via oversampling

## Acknowledgements

The authors acknowledge support from Pablo Tamayo on the many helpful discussions that made this work possible. The authors also acknowledge funding support from the Cancer Therapeutics Training Program from UC San Diego.

## Author Contributions

EB conceived and managed the research. EB led, designed, and conducted analyses. KS, EB, MN, KI, and ST conducted experiments and/or computational analyses. EB, MN and MS oversaw experiments and analyses. EB and KS wrote the manuscript. All authors read and approved of the manuscript.

