## Supplementary Information for "Machine learning of three-dimensional protein structures to predict the functional impacts of genome variation"

### Supplementary Figures and Tables

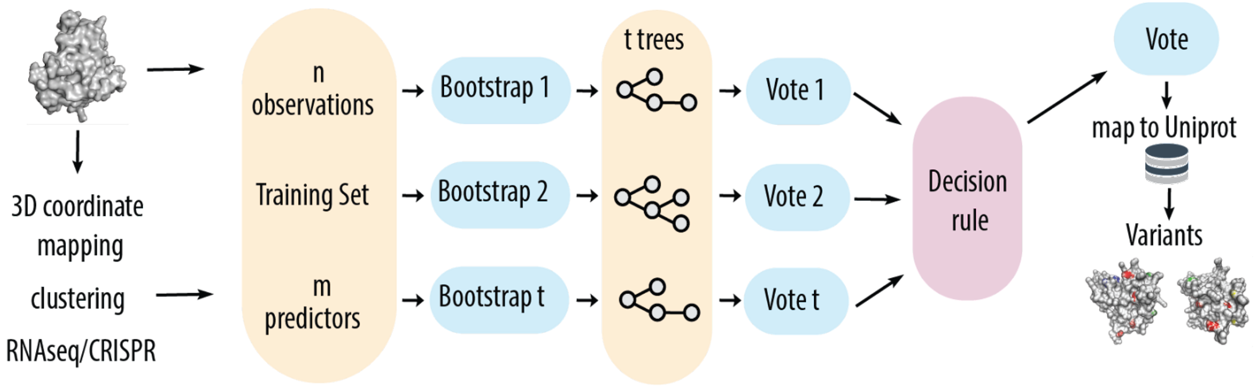


SI1: The Machine Learning workflow used in VAMOS to identify FRVs with respect to the relevant targets. Input features are the 3D mutation coordinates, density based clustering information, and RNAseq expression or CRISPR data. We use a Random Forest Algorithm.

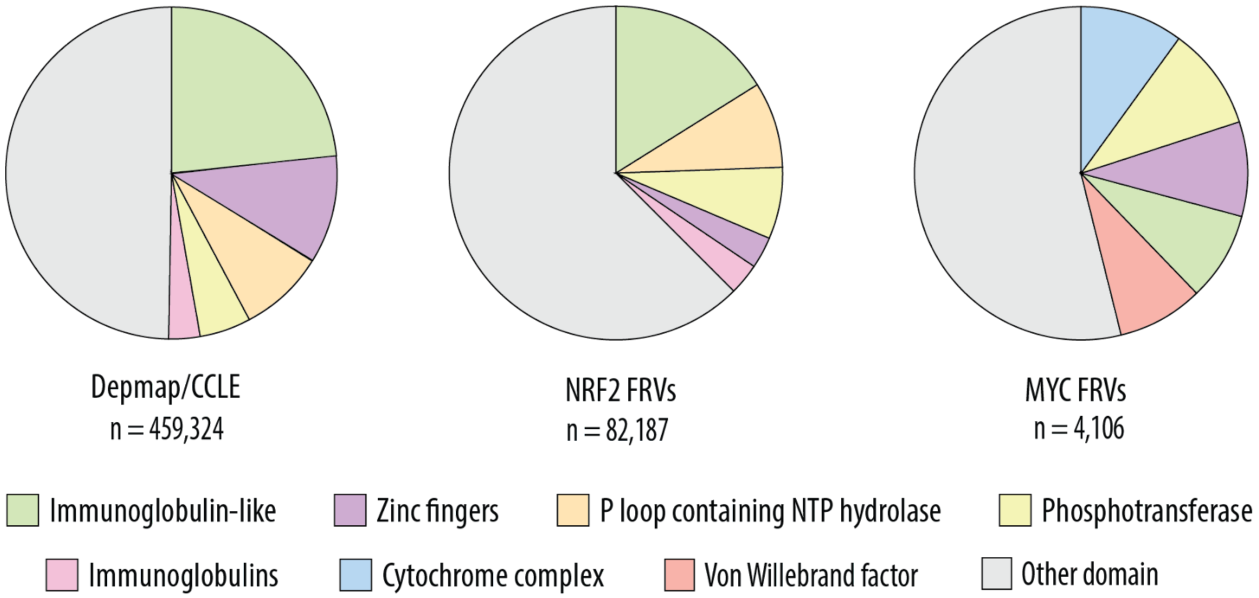


SI2: Top 5 domains present in Depmap/CCLE as a whole, and in the FRVs identified in Depmap/CCLE with NRF2 and MYC as targets. Domain information was obtained from CATH DB only for structures which had been experimentally resolved.

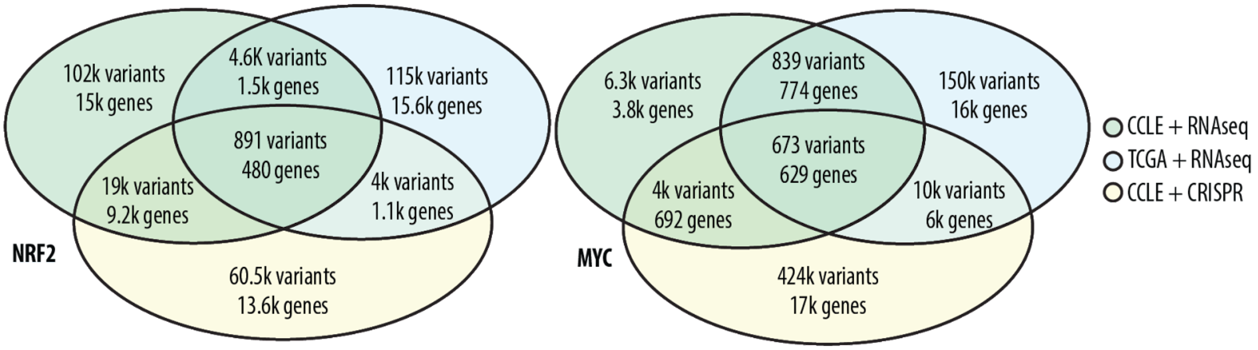


SI3: Pre filtering breakdown of Class 1 mutations across the three datasets

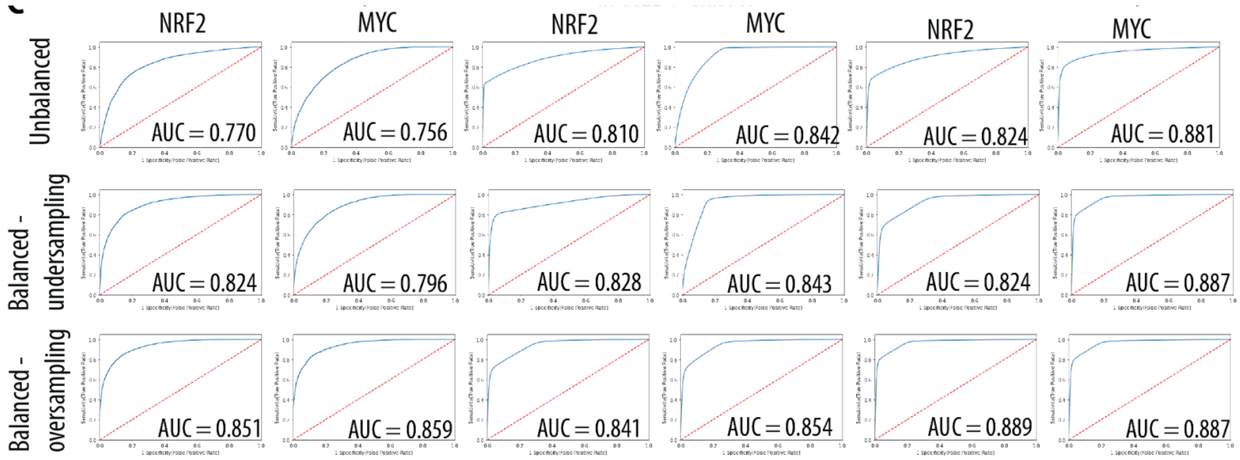


SI4: Additional ROC plots for different sampling conditions

SI5: Scoring metrics for the machine learning algorithm applied to the test data set

| | Nrf2 | | | c-MYC | | |
|---|---|---|---|---|---|---|
| | Cell lines + RNAseq | Cell lines + CRISPR | Tumors + RNAseq | Cell lines + RNAseq | Cell lines + CRISPR | Tumors + RNAseq |
| Accuracy | 0.865 ±0.007 | 0.930 ±0.005 | 0.900 ±0.005 | 0.962 ±0.004 | 0.947 ±0.003 | 0.832 ±0.008 |
| Precision | 0.793 ±0.009 | 0.881 ±0.008 | 0.839 ±0.007 | 0.928 ±0.006 | 0.972 ±0.004 | 0.753 ±0.009 |
| Recall | 0.995 ±0.001 | 0.998 ±0.001 | 0.997 ±0.001 | 0.998 ±0.001 | 0.921 ±0.001 | 0.995 ±0.001 |
| F1 score | 0.883 ±0.005 | 0.935 ±0.005 | 0.911 ±0.004 | 0.962 ±0.003 | 0.946 ±0.003 | 0.858 ±0.006 |
| MCC score | 0.755 ±0.012 | 0.866 ±0.010 | 0.815 ±0.009 | 0.927 ±0.007 | 0.888 ±0.010 | 0.998 $\pm 5.04 * 10^{-6}$ |

*standard deviation values are from 5-fold cross validation

*balanced via oversampling
